# HAHmiR.DB: A Server Platform For High Altitude Human miRNA-Gene Coregulatory Networks And Associated Regulatory-Circuits

**DOI:** 10.1101/2020.05.20.106872

**Authors:** Apoorv Gupta, Ragumani Sugadev, Yogendra Kumar Sharma, Bhuvnesh Kumar, Pankaj Khurana

## Abstract

Rapid ascent to High Altitude (HA) can cause severe damage to body organs and may lead to many fatal disorders. During induction to HA, human body undergoes various physiological, biochemical, hematological and molecular changes to adapt to the extreme environmental conditions. Many literature references hint that gene-expression-regulation and regulatory molecules like microRNAs (miRNAs) and Transcription Factors (TFs) control adaptive responses during HA-stress. These biomolecules are known to interact in a complex combinatorial manner to fine-tune the gene expression and help in controlling the molecular responses during this stress and ultimately help in acclimatization. HAHmiR.DB (High-Altitude Human miRNA Database) is a unique, comprehensive, curated collection of miRNAs that have been experimentally validated to be associated with HA-stress; their level of expression in different altitudes, fold change, experiment duration, biomarker association, disease and drug association, tissue-specific expression level, Gene Ontology (GO) and Kyoto Encyclopaedia of Gene and Genomes (KEGG) pathway associations. As a server platform it also uniquely constructs and analyses interactive miRNA-TF-Gene coregulatory networks and extracts regulatory-circuits/Feed Forward Loops (FFLs) using in-house scripts. These regulatory circuits help to offer mechanistic insights in complex regulatory mechanisms during HA stress. The server can also build these regulatory networks between two and more miRNAs of the database and also identify the regulatory-circuits from this network. Hence HAHmiR.DB is the first-of its-kind database in HA research which a reliable platform to explore, compare, analyse and retrieve miRNAs associated with HA stress, their coregulatory networks and FFL regulatory circuits. HAHmiR.DB is freely accessible at http://www.hahmirdb.in

## INTRODUCTION

High Altitude (HA) is defined as height between 2500 – 4000 m above sea level; very high altitude is between 4000 – 5500 m and altitudes above 5500 m are considered extremely high altitude. These altitudes affect normal physiology and health due to the low partial pressure of oxygen at these altitudes (1). Oxygen concentration and the barometric pressure at sea level is about 21% and 760 mmHg respectively but the barometric pressure decreases gradually and significantly at higher altitudes which also affects the oxygen concentration. At an altitude of 3500m, there are 40% fewer oxygen molecules per breath (1). Hence the supply of oxygen to the body tissues also decreases significantly. This condition is known as hypobaric hypoxia (2). To oxygenate the body adequately, the hypoxic condition triggers an acclimatization mechanism both at the physiological and molecular levels. At the physiological level, the body hyperventilates to increase the oxygen concentration in the blood. The blood pressure increases to augment the oxygen supply to the tissues (3). But the sudden increase in blood flow can also cause fluid to leak from the blood capillaries and this fluid build-up in the lungs and the brain, triggers life-threatening illnesses i.e. High-Altitude Pulmonary Edema (HAPE) and High-Altitude Cerebral Edema (HACE)(4). At the molecular level, the lack of oxygen (hypoxia) is sensed by the oxygen sensor of cells called Prolyl hydroxylases (PHDs)(3). These PHDs further activate the master transcription factor of hypoxia known as hypoxia-inducible factor‐1 (HIF‐1) (5) which largely controls the signalling machinery responsible for hypoxia-adaptive-responses in the body (3). These molecular responses in hypoxia-adaptation are controlled by the fine-tuning of the transcriptome expression(6). The gene regulatory elements, such as TFs and miRNAs are crucial as they have been found to play a decisive role in maintaining physiological and molecular homeostasis, both under normoxic and hypobaric hypoxia conditions(7). miRNAs are known to regulate a number of molecular mechanism during HA acclimatization or disorders, e.g. miR-16, −20b, −22 and −206 and 17/92 are downregulated during HAPE and are responsible for disruption of the ion channels and loss of cellular integrity (8). Increase in red blood cell count, haematocrit values and haemoglobin are established physiological response at HA. Hsa-miR-210-3p (also known as the master regulator hypoxiamiR) expression is positively correlated with the change in red blood cell counts, haemoglobin and haematocrit values at HA and is proposed to be associated with human acclimatization to life at HA (9). Human plasma miRNA expression profiles are thus reported to be associated with human adaptation to hypobaric hypoxic environments (10). hsa-miR-369-3p, hsa-miR-449b-3p, and hsa-miR-136-3p, miR-495 and miR-323a-3p, miR-500a-5p and miR-501-5p have been reported to be differentially expressed at different altitude and duration of exposure and have been proposed to have immense potential as markers or therapeutic targets (11). miRNAs are known to interact with other regulatory molecules like TFs. SNP variants of some of the TFs like EPAS1, EGLN1, PPRAG inherent in the Tibetan population are reported as important ecological traits for high-altitude adaptation (12–14). Several reports hint towards TF and miRNA working in conjunction to influence the precise control and fine-tuning of gene expression during multifactorial disorders like myocardial infarct, cancer, multiple sclerosis, schizophrenia etc. They are also known to regulate a variety of processes in hypoxia associated disorders like malignancies, wound repair, stroke etc. (15–20).

A tripartite interaction between a miRNA, TF and a common Gene form regulatory-circuits also known as a Feed Forward Loop (FFL). In FFL, either the TF or miRNA or both regulate each other and also a target gene. It can be further categorized as miRNA-FFL and TF-FFL. In a miRNA-FFL, miRNA is the main regulator, which regulates a TF and its common target gene while in a TF-FFL, TF is the main regulator (20). These FFLs motifs have been found to play vital roles in disease pathology, drug repurposing, and disease recurrence and hence can be used for the understanding of the underlying mechanism of disease initiation, progression, and recurrence (16,21). Recently, miRNA based FFLs are proposed as potential biomarkers for complex multifactorial disorders like colorectal cancer, myocardial infarct etc. (21,22). Analysis of miRNA-TF-gene co-regulatory networks in these diseases has found to successfully predict the disease pathology and recurrence.

Recently there have been number of studies assessing miRNA expression and regulation during HA stress. This molecular data is scattered and our study is a systematic attempt to collect, curate, analyse, visualize and store this data. HAHmiR.DB (High Altitude Human miRNA Database) is a first-ever comprehensive database for miRNA associated with HA-stress; their coregulatory networks and regulatory-circuits. The database currently contains 386 human miRNAs which are manually curated from peer-reviewed publications related to high-throughput techniques such as microarray, qPCR, RNA seq, etc. The database stores the association of each miRNA with HA-stress in terms of the experimental group, altitude and duration of experiment, level of regulation (up/down), fold change, GEO accession, association as biomarker and a corresponding link to the respective publication. The database also collates mature miRNA Ids, miRNA and RNAcentral accession, family, precursor and mature sequence and the stem-loop structure. It also retrieves and stores the miRNA gene targets. Using these, miRNA-TF-gene co-regulatory networks are built and two types of FFLs i.e. miRNA-FFL and TF-FFL are extracted using in-house scripts. The database also stores other information about miRNAs like their role in other diseases, its tissue-specific expression and their association with drugs. GO and KEGG pathway enrichment of the miRNA targets are also stored. As a server platform, HAHmiR.DB also builds interactive and dynamic miRNA-TF-gene co-regulatory networks of user-defined miRNA list and also identifies the FFL regulatory-circuits from the network. HAHmiR.DB will help to uncover the underlying cross-talk between miRNA, TF and gene that governs the fine-tuning of gene expression during HA acclimatization. It will be useful in revealing novel and robust targets and important regulating modules responsible for acclimatizing to HA. Researchers can further use this information for getting mechanistic insights in the complex molecular responses in HA-related disorders.

## MATERIAL AND METHODS

### Data Collection

The combination of keywords such as “High Altitude” and “miRNA” were used to extract the list of publications from PubMed and Google scholar as on January 2020 (23,24). After removing redundancy and duplicity, the publications were manually curated to identify the differentially expressed (DE) miRNAs from human studies. For each miRNA its experimental group, experiment altitude, duration of induction, level of expression, fold Change, GEO (Gene Expression Omnibus) accession and its reference paper were compiled. Sometimes the information e.g. fold change, the tissue of expression was not categorically listed in the publication, so those entries are recorded as ‘na’. We also recorded whether any literature text had associated the miRNA as a biomarker for HA acclimatization. Subsequently, miRBase stem Loop Id, miRBase accession number, miRNA Family, 5’ ID, 3’ID, Chromosome number, precursor sequence, mature sequence, and RNA central ID for each miRNAs were retrieved and stored from the miRBase(25), miRGen 2.0(26) and RNAcentral(27) databases respectively. To ensure uniformity in the nomenclature, the names of precursor miRNAs were mapped to the mature miRNA using miRBase(25) and miRDB(28) databases.

### Data Annotation

A comprehensive and exhaustive list of human TF and Co-Transcription Factors (Co-TFs) were mined from TFcheckpoint, DBD, TcoF-DB V2 and TRANSFAC(29–32). This was stored as human TF list. For each miRNA, its experimentally-validated gene targets were identified from MirTarBase and miRecords (33,34). These gene targets were annotated as TF or gene by comparison with human TF list. This way the corresponding miRNA→gene interactions were annotated as miRNA→TF or miRNA→gene interactions.

Further TF → gene and TF → miRNA interactions were compiled from OregAnno(35), TRRUST-V2(36) and TransmiR v2.0(37), PutmiR(38) databases respectively. Using these miRNA→gene, miRNA→TF, TF → gene, TF → miRNA interactions, a miRNA-TF-gene coregulatory network was built for each miRNA in the database. Additionally miRNA → miRNA interactions between miRNAs in the database were also added from PmmR(39) database to identify more complex network interactions.

To make the database more informative, several other attributes were added; miRNA: disease associations were mined from HMDD(40); miRNA: tissue association from miRmine(41) and miRNA:drug relationship from PharmacomiR(42). For each miRNA, GO-functional enrichment and KEGG pathway enrichment of their target genes was performed using Database for Annotation, Visualization and Integrated Discovery (DAVID)(43). The complete overview of data collection and annotation is listed in (Figure 1)

**Figure 1.**
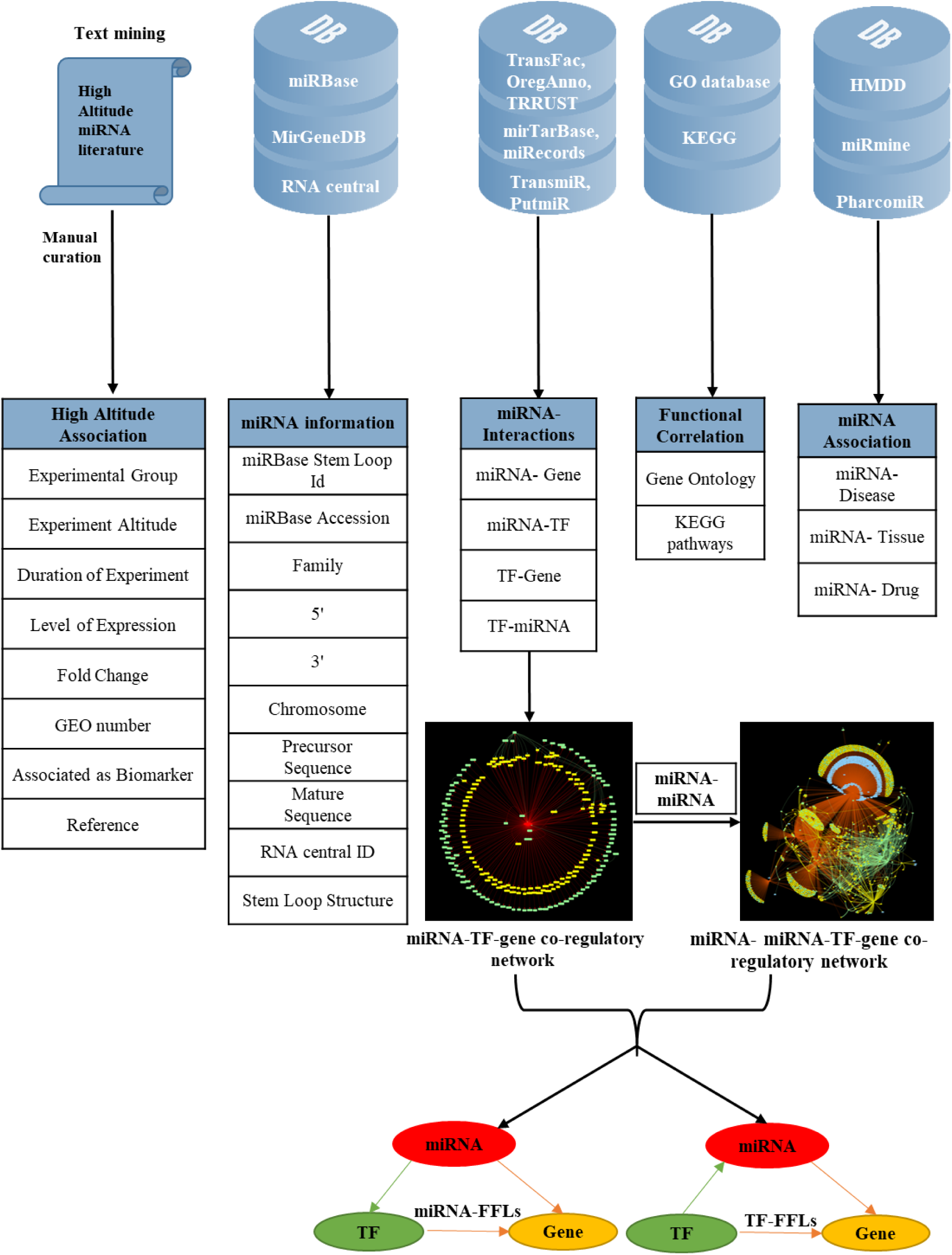
Overview of data collection and annotation in HAHmiIR.DB

### FFL Motif Identification

Each vertex in the network is labelled as V_m_, V_TF_ or V_gene_. Where, V_m_ refers to a miRNA, V_TF_ refers to a TF and V_gene_ refers to a gene. The edges are annotated as E_mt_, E_mg_, E_tg_, E_tm_ where E_mt_ refers to edge from V_m_ to V_TF_, Emg refers to edge from V_m_ to V_gene_, E_tg_ refers to edge from V_TF_ to V_gene_ and E_tm_ refers to edge from V_TF_ to V_m_.

For each V_m_, its edges E_mg_ from the miRNA→gene interactions is searched in miRNA→TF interaction to identify a corresponding Emt with a common vertex Vm. Thereafter, an analogous Etg is searched in the TF → gene interactions. If found, the complete graph containing three vertices (V_m_, V_TF_, V_gene_) and edges(E_mt_, E_mg_, E_tg_) were labelled as miRNA-FFL motif graph.

Similarly, in order to find TF-FFL motif, edge E_tm_ and E_tg_ were identified from TF → miRNA and TF → gene interactions dataset respectively. Further the program scans for an E_mg_ associated with V_m_ and V_gene_ in miRNA→gene interaction dataset. If algorithm finds E_mg_, the complete graph containing three vertices (V_m_, V_TF_, V_gene_) and edges (E_tm_, E_mg_, E_tg_) were labelled as TF-FFL motif graph. This methodology was used to identify miRNA-FFLs and TF-FFLs for each human miRNA.

### Database Development

Finally, after collection, processing and enrichment of the data, all the database files were stored as JavaScript Object Notation (JSON) files in MongoDB database relational database(44). These JSON files was uploaded on the server localhost using pymongo and query commands were made in the command line client in MongoDB compass. The database uses Asynchronous JavaScript and XML (AJAX) technique for API calls. AJAX is a web development technique that is used for creating interactive web applications. It utilizes XHTML for content along with the document object model and JavaScript for dynamic content display. Vis.js library functions are used for interactive visualization of network graphs on front-end. The website is available online at www.hahmirdb.in and requires no registration. It provides a user-friendly interface to browse and analyse HA associated miRNAs.

## RESULTS

HAHmiR.DB database allows to browse, retrieve, analyse and compare miRNAs which are associated with HA-stress. In ‘Browse’ option, user can either search individual miRNA directly from a pull-down menu or can use a variety of filters to fetch the miRNAs of interest. The “Search by Single miRNA” option allows the user to select a miRNA from a drop-down menu and further link to its detailed information page. The database also provides user with four other filters to select and retrieve the list of miRNAs (Figure 2). These filters are based on level of expression (upregulated/ downregulated); duration of experiment (Days/Months/Native), experimental altitude (3600-7000m) and by search by association. The experimental altitude option is provided with a slider bar, where user can select the altitude range (3600m - 7600m) to get the set of miRNAs associated with that extent. The fourth browse option of the database “Browse by Association” is the special option, where user can retrieve a list of miRNAs associated with gene list/biomarker/drug/GO ID/GO term /KEGG ID/KEGG term.

**Figure 2:**
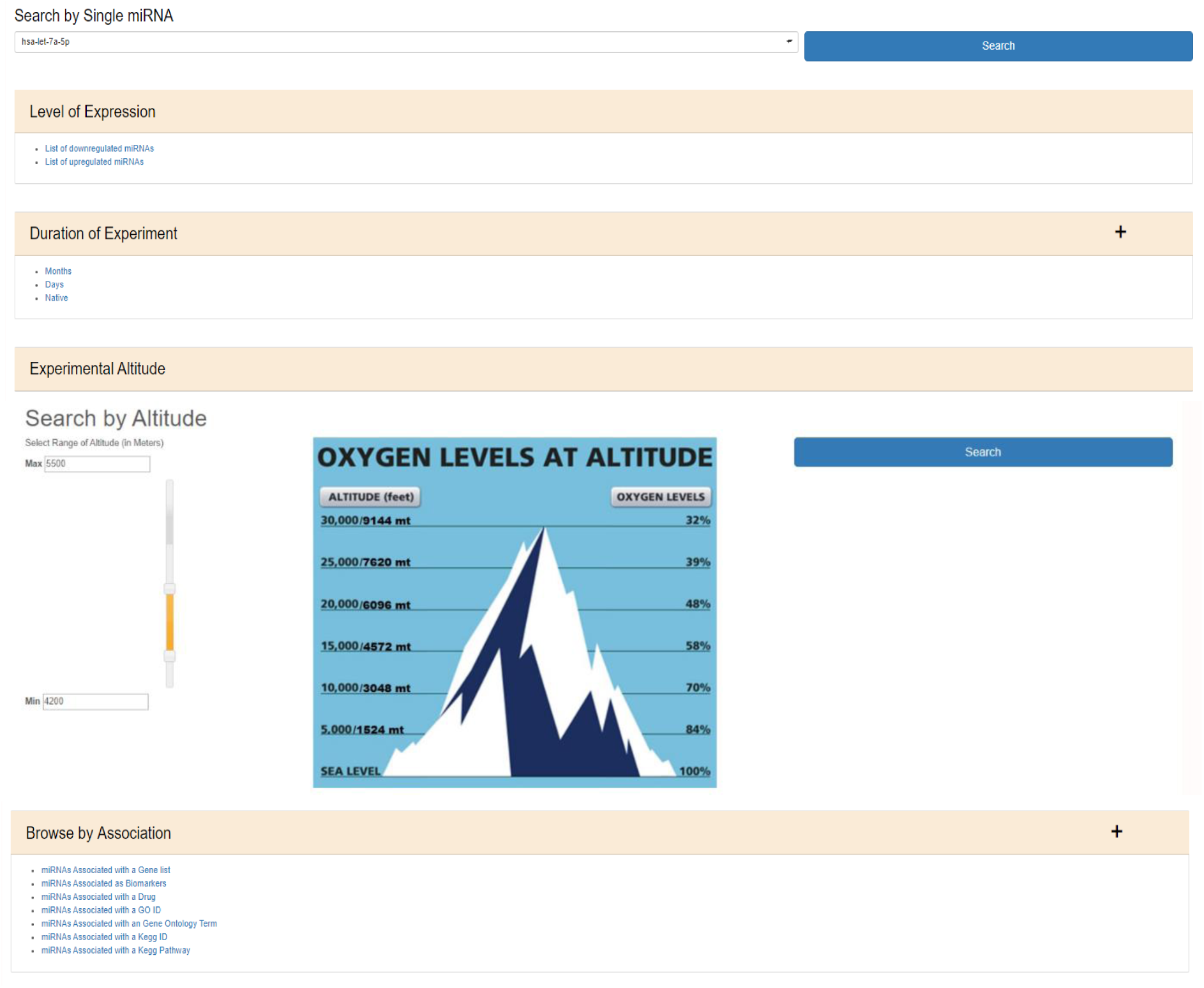
The web image of HAHmiR.DB - ‘Browse’ option allows the user to browse DE miRNA during HA using multiple filters i.e. level of expression, duration of experiment, experimental altitude and browse by associations that includes filters such as miRNAs associated with gene list, miRNAs associated as biomarker, miRNAs associated with a drug, miRNAs associated with an GO ID/term, miRNAs associated with a KEGG ID/pathway

“miRNAs Associated with a Gene List” option allows to select gene(s) from a pull-down gene list and fetch the miRNAs regulating these genes (Supplementary Figure S1a). “miRNAs Associated as Biomarkers” option allows the user to extract list of miRNAs that are associated as biomarkers in HA conditions (Supplementary Figure S1b). Similarly, “miRNAs Associated with a Drug” option allows user to identify miRNAs associated with a particular drug (Supplementary Figure S1c). “miRNAs Associated with a GO ID” (Supplementary Figure S1d), “miRNAs Associated with a GO Term” (Supplementary Figure S1e), “miRNAs Associated with a KEGG ID” (Supplementary Figure S1f), “miRNAs Associated with a KEGG Pathway” (Supplementary Figure S1g) allows user to fetch miRNA based on functional association such as GO ID, GO term, KEGG ID and KEGG pathway respectively. All the above options provide a resultant list of miRNAs that are hyperlinked to their respective detailed information pages. The list of these miRNAs can be downloaded in Excel /PDF format for further analysis.

The miRNA information page of HAHmiR.DB can be broadly divided into four sections.

### (i) Knowledge base

The first section of the database provides general information about miRNA such as miRBase accession number, miRBase Stem Loop Id, 5’/3’ ID, miRNA family, chromosome number, precursor and mature sequences, RNAcentral ID etc. It also contains miRNA stem loop structure where mature miRNA sequence is highlighted with red colour (Figure 3a). The miRBase accession, 5’/3’IDs and RNA Central Ids are also hyperlinked to miRBase and RNAcentral databases to provide additional details of the miRNA like mature and precursor miRNA sequence, miRNA stem loop structure, chromosome location, source organism, target proteins, target lncRNAs, genome locations, Rfam classification etc (25,27).

**Figure 3:**
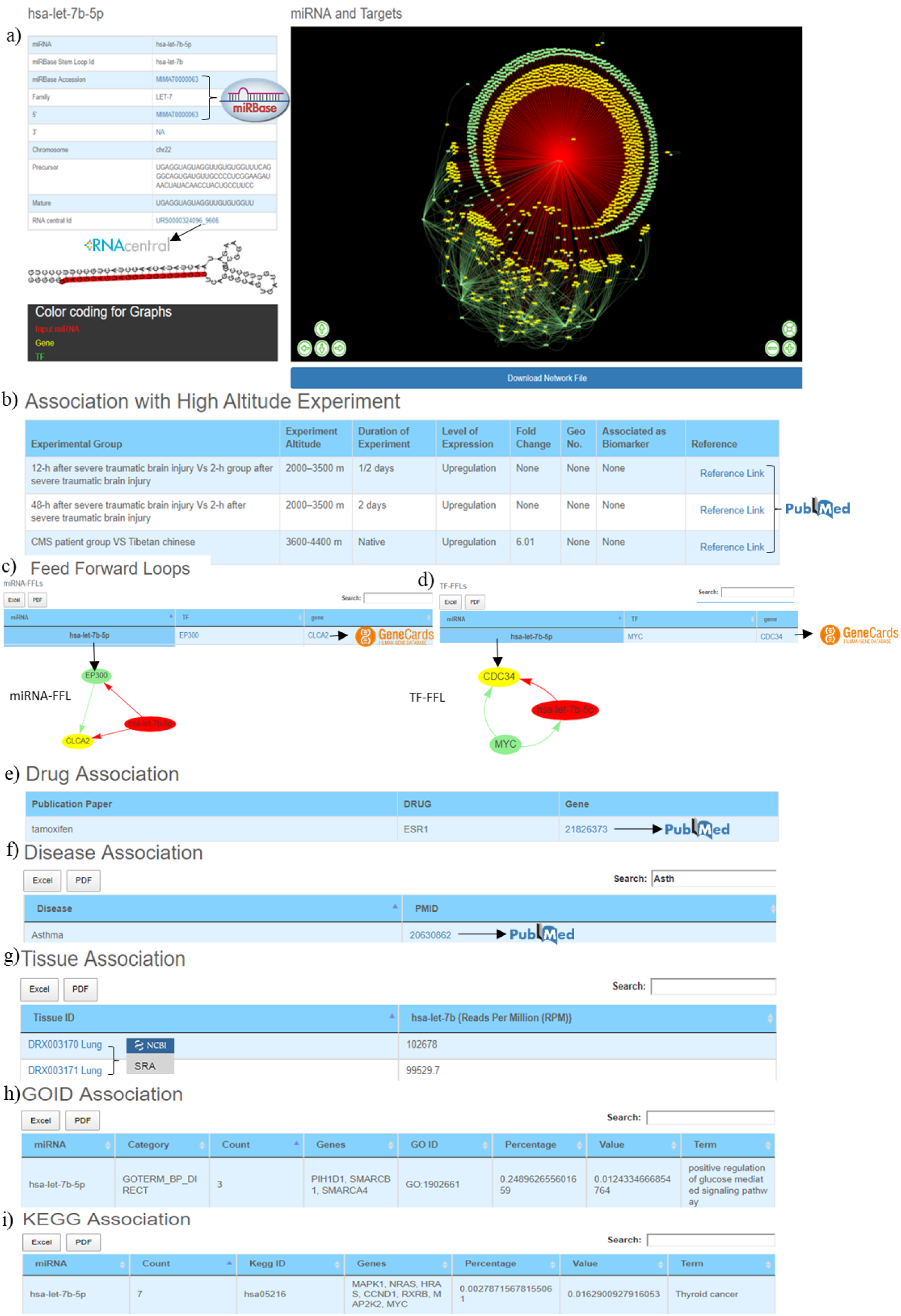
Web image of the miRNA information page a) The first section of hsa-let-7b-5p miRNA information page with details about the miRNA and hyperlinking to external databases like miRBase and RNAcentral b) The second section shows information related to experimental details of DE human miRNA during HA conditions in published literature. c) The web image of the miRNA-FFL in tabular format. d) The web image of the TF-FFL in tabular format. e) miRNA associations with drug f) miRNA associations with different disease g) miRNA expression level in different tissues. h) GO Functional enrichment of the miRNA targets i) KEGG pathway enrichment of the miRNA targets. The lengthy tables can be searched through ‘search’ box and information in table can be download in both excel and PDF format

HAHmiR.DB uniquely offers directed miRNA-TF-gene co-regulatory network of the miRNA. The details of the in-house scripts to build this network are provided in the methodology section. The network nodes are color coded (miRNA-red, TF-green, Target Gene-yellow) and interactive with option likes Zoom in/out, translation of nodes etc. These networks can be saved in publication quality images. User can also download the network file which can also be exported and visualized in publically freely available software’s like Cytoscape(45), BINA(46), Gephi etc.

### (ii) miRNA association with high altitude

For each miRNA, its association with HA is compiled in the form of experimental group, experiment altitude, duration of experiment, level of expression, fold change, GEO accession, association as biomarker and the respective reference which is hyperlinked to PubMed (23) database which allows easy access to the original publication (Figure 3b).

### (iii) Feed Forward Loops

Biological systems are often built of regulatory-circuits that carry out key functions (47). These recurring patterns are very common in Gene Regulatory Networks (GRNs) and are known as network motifs. Important regulatory molecules such as miRNA and TFs often follows these recurring pattern in co-regulatory networks to control the complex molecular and cellular responses of living cells (48). FFLs are most overrepresented motifs that are found in the co-regulatory networks (49). They are of two types miRNA-FFLs and TF-FFLs based on the interaction between miRNA and TF. If the interaction is of the type that a miRNA dysregulates TF and both together regulate target gene than it is called a miRNA-FFL. A type where a TF regulates a miRNA and both then regulate a target gene, then its known as TF-FFL.

Both these regulatory-circuits/FFLs are extracted from miRNA-TF-gene coregulatory networks using in-house scripts as discussed in the methodology. The FFLs are provided in a tabular format (Figure 3c, 3d) which can be explored and downloaded. Clicking on the miRNA would provide a network of the FFL in a pop-up window. Each TF and TG symbol in the FFL table are further hyperlinked to the GeneCards(50) database. This would serve as a ready-reference for the user to get additional details about the gene that includes aliases of genes, promoter and enhancer location of gene, protein coded by gene, functional characterization, cellular localization, pathway enrichment, gene-gene interaction network, drug-gene relationship, tissue specific gene expression profile, orthologs, paralogs, transcript variant etc. These regulatory-circuits/FFLs could offer mechanistic insights during complex cellular responses and also open new horizons in HA research for identifying potential candidate markers for HA acclimatization.

### (iv) Association of miRNA with Drugs, diseases and Tissues

This section provides details of the drug, disease and tissue association of the miRNAs. The information is represented in three different tables. The first table shows information about miRNA and its associated drug compiled from PharmocomiR (Figure 3e). A direct link to the corresponding reference in PubMed is provided for ready reference. This information may help the user to design/validate miRNA-based drug-targeting/repurposing experiments. Similarly, miRNA-disease-associations (Figure 3f) and miRNA tissue-specific-expression (Figure 3g) are also provided in tabular formats. Each entry in tissue specific expression table have been hyperlinked to the SRA database(51), which provides additional tissue-specific experimental details (Figure 3g).This could be important to design experiments which aim to study tissue-specific expression. These tables are equipped with the integrated “search” option and tables can also be downloaded in Excel/PDF format.

#### Association with GO and KEGG pathways

This section presents the functional characterization of the miRNA gene targets. Both functional and pathway enrichment are stored and visualized in a tabular format. These tables are provided with a “search” option that allows exploring the extensive tables and fetching the user-defined information (Gene/GO-term/KEGG pathway) easily. Figure 3h shows the GO table searched with “calcium” keyword to identify specific GO term from the 203 GO entries of hsa-let-7b-5p. Similarly Figure 3i the “PI3K” keyword was used to search specific entry from KEGG pathway entries of hsa-let-7b-5p.

HAHmiR.DB server also allows user to perform integrated network analysis through “Explore” option of the database. In this option, user can select multiple (upto maximum of 4) miRNAs from the database. The server then builds a dynamic miRNA-TF-gene co-regulatory network between these miRNAs. The network include miRNAs, their targeted TFs, targeted genes, and also its interacting miRNAs. The network is presented as an interactive visualization with color coding and zoom-in and translation options. (Figure 4). The network image may be saved as publication quality images and can also be downloaded. This file can then be used to visualize networks offline in visualization software like cytoscape(45), BINA(46), Gephi etc. HAHmiR.DB also extracts the FFLs in the complex network and presents them in a tabular format. User can click on each miRNA in the row to see its tripartite graph as pop-ups. The genes and TFs are hyperlinked to GeneCards database. The integrated search option is provided with the table for searching the FFLs using TF/Gene. The FFL table can be downloaded in excel and PDF format.

**Figure 4:**
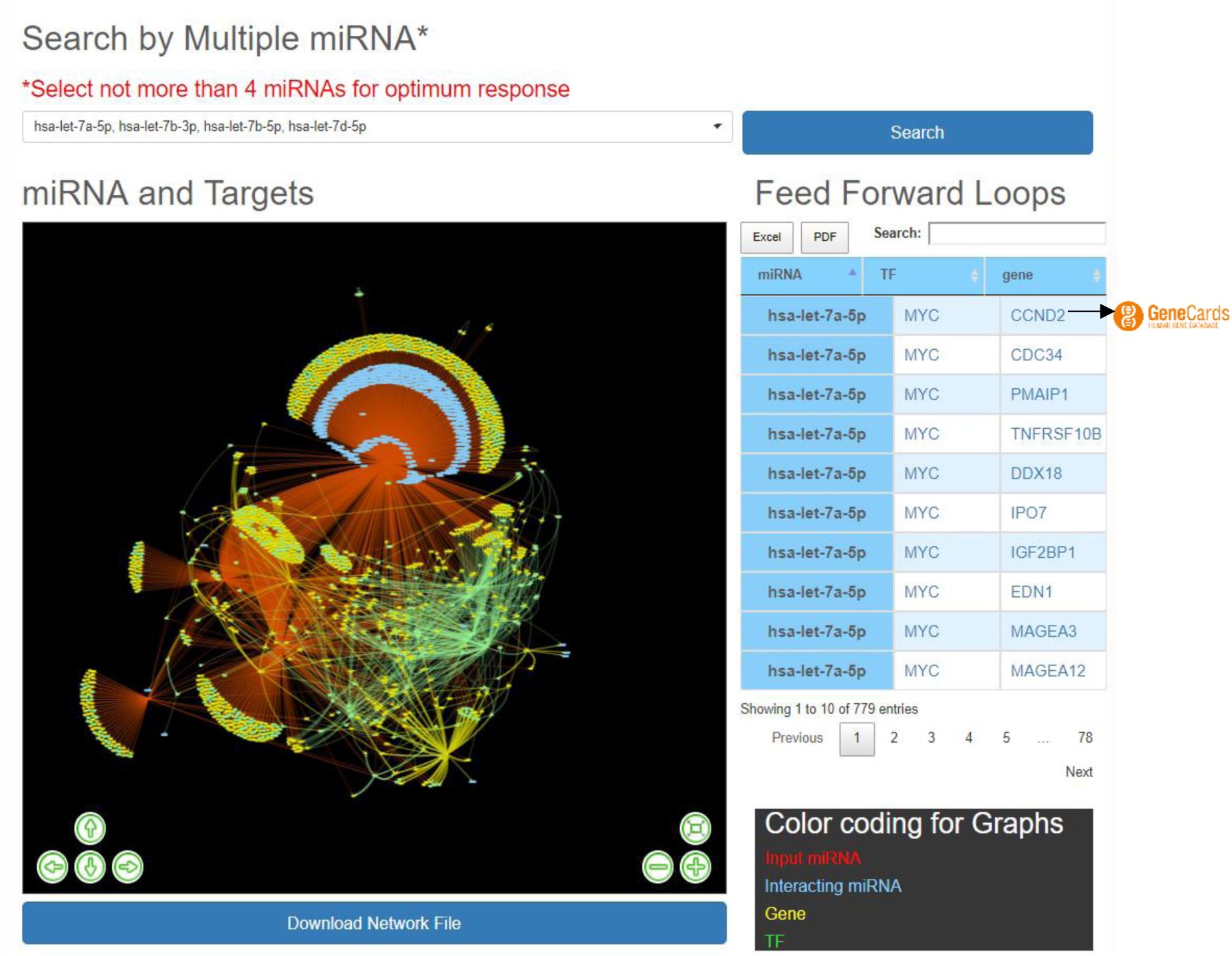
“Explore” option of the database. The web image of the section shows complex network of four miRNA i.e hsa-let-7b-5p, hsa-let-7a-5p, hsa-let-7b-3p, hsa-let-7d-5p and identified FFLs in the adjacent table.

#### HAHmiR.DB Statistics

HAHmiR.DB contains 556 associations of 386 miRNAs that are differentially expressed at HA. In terms of interactions HAHmiR.DB consist of ~56,000 miRNA target interactions which can be further divided into 38,496 miRNA→ gene, 17,974 miRNA→ TF interactions. Additionally the database contains 47,115 TF→ gene and 1500 miRNA→ TF interactions. The database also contains 449 unique miRNA-disease associations and 77 miRNA-drug associations. In the database, 52% miRNAs are downregulated and 48% are upregulated. The 556 miRNA entries can also be divided into three categories based on the duration of experiment, 43% of studies had the duration of experiment ranging in months, 40% of studies ranging in days and rest 17% were studies related to HA natives.

For the genes in the database, two different types of functional characterizations were performed i.e GO and KEGG pathway enrichment. The GO enrichment shows the ‘mitotic cell cycle regulation’, ‘regulation of apoptosis process’, ‘cellular response to oxygen levels’, etc. as the top biological processes (Figure 5a). ‘Cell cycle regulation’ is an important cellular mechanism induced by the hypoxic stress that governs the cell fate from cell proliferation to apoptosis and is widely reported and studied biological process during hypoxic environments (52). The ‘cellular response to oxygen level’ is governed by Prolylhydroxylases (PHD) oxygen sensors of cells that controls the activation of HIF during low oxygen condition which further regulates hypoxic stress adaptative responses (7,52–54). Similarly ‘transcription factor binding’, ‘protein kinase binding’, ‘cell adhesion molecule binding’, etc. and ‘nucleoplasm (nucleus & cytoplasm)’, ‘nucleolus’, etc. are the top molecular functions and cellular components respectively (Figure 5b and 5c). These pathways have been reported as hallmark responses to HA stress. Literature shows that protein kinases present in both nucleus and cytoplasm are signal transducers that play important roles in transducing hypoxic stress signals from the cytoplasm to the nucleus for the activation of hypoxic stress-responsive transcription factors like HIF1, NFKB1 p53 etc(52). Also the downregulation of cell adhesion proteins like PECAM-1, ZO-1 during hypobaric hypoxia has been correlated as a signal for vascular leakage and (High Altitude Pulmonary Edema) HAPE(55).

**Figure 5:**
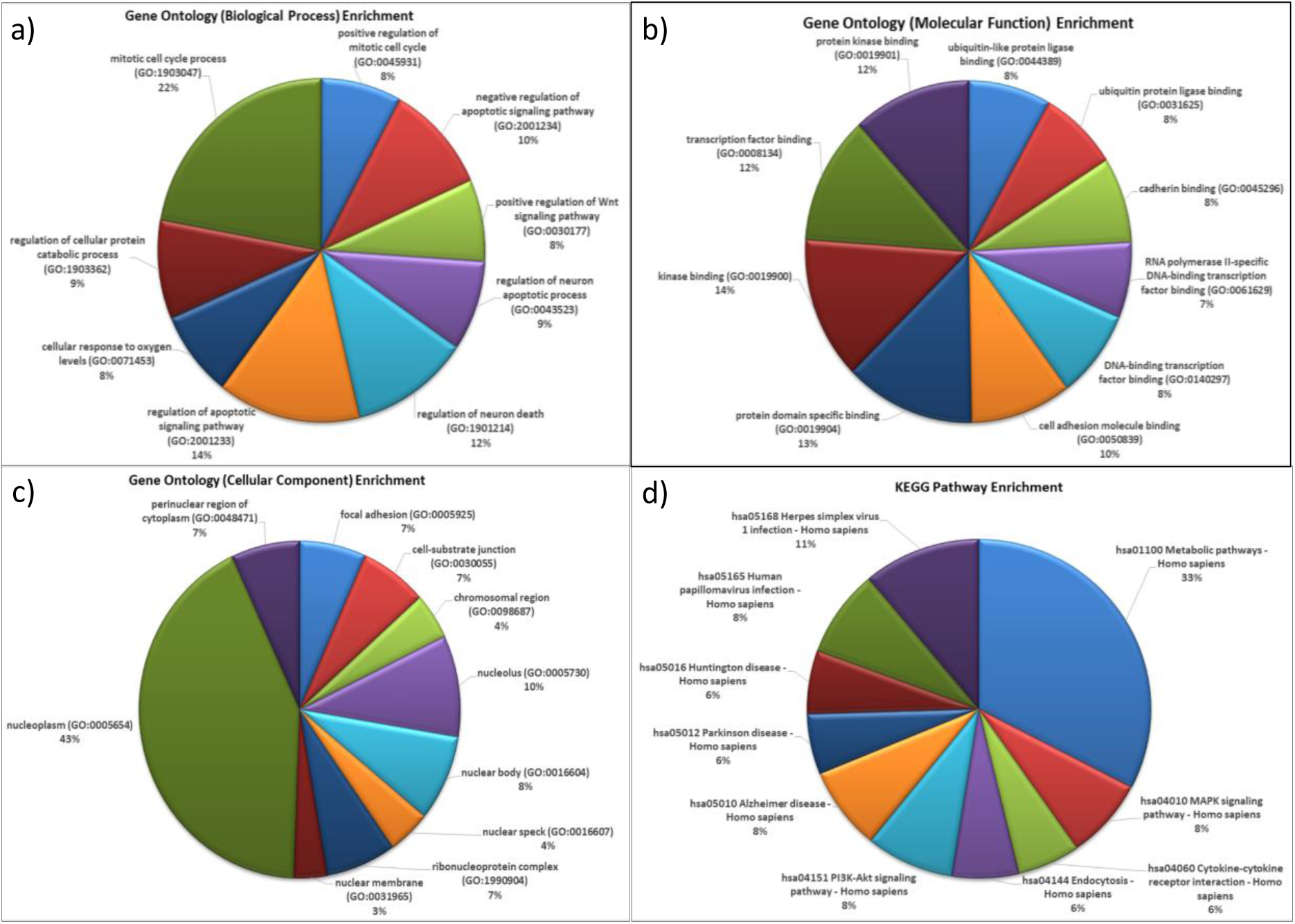
Functional characterization of differentially expressed HA miRNA-targets a) GO: Biological processes. b) GO: Molecular functions c) GO: Cellular Compartment and d) KEGG Pathway Enrichment.

The KEGG pathway enrichment shows ‘metabolic pathway’ as most enriched in the miRNA-targets (Figure 5d). Metabolic pathways are responsible in maintaining body-weight homeostasis during HA adaptation/acclimatization (56).

The functional annotation of miRNAs target genes in database shows correlations of biological processes, molecular functions and pathways with hallmark responses to hypobaric-hypoxic stress and other HA stress conditions (7,52–54). Hence this supports the comprehensive biomolecular structure of the database. Thus HAHmiR.DB is a comprehensive, user-friendly repository of human miRNAs, TFs and genes to study HA stress responses at molecular level.

## DISCUSSION

HAHmiR.DB is an interactive resource for HA associated Human miRNAs. The collection has been stored in the MongoDB database that can be updated from time to time. JSON helps in easy and smooth transferring of the data between the server and API. Currently, there is only a single database for HA species genome dataset “The Yak Genome Database” that provides information on genomic sequence related of YAK (HA resident animal species) (57). HAHmiR.DB is the only database that has a collection of manually curated DE high altitude specific human miRNAs that are fetched and manually curated from the literature. The expression profile of a miRNA from different publications are compiled at one place so that it can be compared, analysed and retrieved at ease. The portal enables the user to browse miRNAs individually or allows batch retrieval based on different query filters. It also provides information about the association of these miRNAs in other diseases, its tissue-specific expression and its pharmacological relation with other drugs. It identifies miRNA-target interactions (MTIs) of each miRNA constructs directed tripartite miRNA-TF-gene co-regulatory network. Subsequently, HAHmiR.DB identifies regulatory-circuits/FFLs in each miRNA-TF-gene coregulatory network using in-house scripts. The database also performs integrative analysis with user-defined miRNAs and constructs directed miRNA-TF-gene coregulatory network of input miRNAs. These miRNA-TF-gene co-regulatory networks contain regular interaction miRNA-gene, miRNA-TF, TF-gene and TF-miRNA with additional miRNA-miRNA interactions. This complex network would help understand the biological mechanisms underlining a molecular trait during HA ascent. The database is the first database that provides miRNA-TF-gene co-regulatory networks and its regulatory-circuits in a pathophysiological stress condition.

## REFERENCES

1. Peacock, A.J. (1998) ABC of oxygen: oxygen at high altitude. Bmj, 317, 1063–1066.

2. Irarrazaval, S., Allard, C., Campodonico, J., et al. (2017) Oxidative Stress in Acute Hypobaric Hypoxia. High altitude medicine & biology, 18, 128–134.

3. Giaccia, A.J., Simon, M.C., Johnson, R. (2004) The biology of hypoxia: the role of oxygen sensing in development, normal function, and disease. Genes & development, 18, 2183–2194.

4. Mehta, S.R., Chawla, A., Kashyap, A.S. (2008) Acute Mountain Sickness, High Altitude Cerebral Oedema, High Altitude Pulmonary Oedema: The Current Concepts. Medical journal, Armed Forces India, 64, 149–153.

5. Bishop, T., Ratcliffe, P.J. (2014) Signaling hypoxia by hypoxia-inducible factor protein hydroxylases: a historical overview and future perspectives. Hypoxia, 2, 197–213.

6. Nakayama, K., Kataoka, N. (2019) Regulation of Gene Expression under Hypoxic Conditions. International journal of molecular sciences, 20.

7. Gupta, A., Sugadev, R., Sharma, Y.K., et al. (2018) Role of miRNAs in hypoxia-related disorders. Journal of biosciences, 43, 739–749.

8. Alam, P., Saini, N., Pasha, M.A. (2015) MicroRNAs: An Apparent Switch for High-Altitude Pulmonary Edema. MicroRNA, 4, 158–167.

9. Yan, Y., Wang, C., Zhou, W., et al. (2016) Elevation of Circulating miR-210-3p in High-Altitude Hypoxic Environment. Frontiers in physiology, 7, 84.

10. Yan, Y., Shi, Y., Wang, C., et al. (2015) Influence of a high-altitude hypoxic environment on human plasma microRNA profiles. Scientific reports, 5, 15156.

11. Norman E Buroker, X.-H.N., Zhao-Nian Zhou, Kui Li, Wei-Jun Cen, Xiu-Feng Wu, Wei-Zhong Zhu, C Ronald Scott and Shi-Han Chen (2013) Circulating miRNAs from dried blood spots are associated with high altitude sickness. Journal of Medical Diagnostic Methods, 2.

12. Peng, Y., Cui, C., He, Y., et al. (2017) Down-Regulation of EPAS1 Transcription and Genetic Adaptation of Tibetans to High-Altitude Hypoxia. Molecular biology and evolution, 34, 818–830.

13. Petousi, N., Robbins, P.A. (2014) Human adaptation to the hypoxia of high altitude: the Tibetan paradigm from the pregenomic to the postgenomic era. Journal of applied physiology, 116, 875–884.

14. Yang, J., Jin, Z.B., Chen, J., et al. (2017) Genetic signatures of high-altitude adaptation in Tibetans. Proceedings of the National Academy of Sciences of the United States of America, 114, 4189–4194.

15. Arora, S., Rana, R., Chhabra, A., et al. (2013) miRNA-transcription factor interactions: a combinatorial regulation of gene expression. Molecular genetics and genomics: MGG, 288, 77–87.

16. Zhang, G., Shi, H., Wang, L., et al. (2015) MicroRNA and transcription factor mediated regulatory network analysis reveals critical regulators and regulatory modules in myocardial infarction. PloS one, 10, e0135339.

17. Hao, S., Huo, S., Du, Z., et al. (2019) MicroRNA-related transcription factor regulatory networks in human colorectal cancer. Medicine, 98, e15158.

18. Li, R., Jiang, S., Li, W., et al. (2019) Exploration of prognosis-related microRNA and transcription factor co-regulatory networks across cancer types. RNA biology, 16, 1010–1021.

19. Guo, A.Y., Sun, J., Jia, P., et al. (2010) A novel microRNA and transcription factor mediated regulatory network in schizophrenia. BMC systems biology, 4, 10.

20. Nuzziello, N., Vilardo, L., Pelucchi, P., et al. (2018) Investigating the Role of MicroRNA and Transcription Factor Co-regulatory Networks in Multiple Sclerosis Pathogenesis. International journal of molecular sciences, 19.

21. McGee, S.R., Tibiche, C., Trifiro, M., et al. (2017) Network Analysis Reveals A Signaling Regulatory Loop in the PIK3CA-mutated Breast Cancer Predicting Survival Outcome. Genomics, proteomics & bioinformatics, 15, 121–129.

22. Lin, Y., Sibanda, V.L., Zhang, H.M., et al. (2015) MiRNA and TF co-regulatory network analysis for the pathology and recurrence of myocardial infarction. Scientific reports, 5, 9653.

23. Topper, L., Boehr, D. (2018) Publishing trends of journals with manuscripts in PubMed Central: changes from 2008–2009 to 2015–2016. Journal of the Medical Library Association: JMLA, 106, 445–454.

24. Falagas, M.E., Pitsouni, E.I., Malietzis, G.A., et al. (2008) Comparison of PubMed, Scopus, Web of Science, and Google Scholar: strengths and weaknesses. FASEB journal: official publication of the Federation of American Societies for Experimental Biology, 22, 338–342.

25. Griffiths-Jones, S., Grocock, R.J., van Dongen, S., et al. (2006) miRBase: microRNA sequences, targets and gene nomenclature. Nucleic acids research, 34, D140–144.

26. Alexiou, P., Vergoulis, T., Gleditzsch, M., et al. (2010) miRGen 2.0: a database of microRNA genomic information and regulation. Nucleic acids research, 38, D137–141.

27. The, R.C., Petrov, A.I., Kay, S.J.E., et al. (2017) RNAcentral: a comprehensive database of non-coding RNA sequences. Nucleic acids research, 45, D128–D134.

28. Wong, N., Wang, X. (2015) miRDB: an online resource for microRNA target prediction and functional annotations. Nucleic acids research, 43, D146–152.

29. Chawla, K., Tripathi, S., Thommesen, L., et al. (2013) TFcheckpoint: a curated compendium of specific DNA-binding RNA polymerase II transcription factors. Bioinformatics, 29, 2519–2520.

30. Kummerfeld, S.K., Teichmann, S.A. (2006) DBD: a transcription factor prediction database. Nucleic acids research, 34, D74–81.

31. Schmeier, S., Alam, T., Essack, M., et al. (2017) TcoF-DB v2: update of the database of human and mouse transcription co-factors and transcription factor interactions. Nucleic acids research, 45, D145–D150.

32. Matys, V., Fricke, E., Geffers, R., et al. (2003) TRANSFAC: transcriptional regulation, from patterns to profiles. Nucleic acids research, 31, 374–378.

33. Xiao, F., Zuo, Z., Cai, G., et al. (2009) miRecords: an integrated resource for microRNA-target interactions. Nucleic acids research, 37, D105–110.

34. Chou, C.H., Shrestha, S., Yang, C.D., et al. (2018) miRTarBase update 2018: a resource for experimentally validated microRNA-target interactions. Nucleic acids research, 46, D296–D302.

35. Griffith, O.L., Montgomery, S.B., Bernier, B., et al. (2008) ORegAnno: an open-access community-driven resource for regulatory annotation. Nucleic acids research, 36, D107–113.

36. Khan, A., Fornes, O., Stigliani, A., et al. (2018) JASPAR 2018: update of the open-access database of transcription factor binding profiles and its web framework. Nucleic acids research, 46, D1284.

37. Tong, Z., Cui, Q., Wang, J., et al. (2019) TransmiR v2.0: an updated transcription factor-microRNA regulation database. Nucleic acids research, 47, D253–D258.

38. Bandyopadhyay, S., Bhattacharyya, M. (2010) PuTmiR: a database for extracting neighboring transcription factors of human microRNAs. BMC bioinformatics, 11, 190.

39. Sengupta, D., Bandyopadhyay, S. (2011) Participation of microRNAs in human interactome: extraction of microRNA-microRNA regulations. Molecular bioSystems, 7, 1966–1973.

40. Huang, Z., Shi, J., Gao, Y., et al. (2019) HMDD v3.0: a database for experimentally supported human microRNA-disease associations. Nucleic acids research, 47, D1013–D1017.

41. Panwar, B., Omenn, G.S., Guan, Y. (2017) miRmine: a database of human miRNA expression profiles. Bioinformatics, 33, 1554–1560.

42. Rukov, J.L., Wilentzik, R., Jaffe, I., et al. (2014) Pharmaco-miR: linking microRNAs and drug effects. Briefings in bioinformatics, 15, 648–659.

43. Dennis, G., Jr., Sherman, B.T., Hosack, D.A., et al. (2003) DAVID: Database for Annotation, Visualization, and Integrated Discovery. Genome biology, 4, P3.

44. Sanchez-de-Madariaga, R., Munoz, A., Castro, A.L., et al. (2018) Executing Complexity-Increasing Queries in Relational (MySQL) and NoSQL (MongoDB and EXist) Size-Growing ISO/EN 13606 Standardized EHR Databases. Journal of visualized experiments: JoVE.

45. Su, G., Morris, J.H., Demchak, B., et al. (2014) Biological network exploration with Cytoscape 3. Current protocols in bioinformatics, 47, 8 13 11–24.

46. Gerasch, A., Faber, D., Kuntzer, J., et al. (2014) BiNA: a visual analytics tool for biological network data. PloS one, 9, e87397.

47. Mangan, S., Alon, U. (2003) Structure and function of the feed-forward loop network motif. Proceedings of the National Academy of Sciences of the United States of America, 100, 11980–11985.

48. Sadeghi, M., Ranjbar, B., Ganjalikhany, M.R., et al. (2016) MicroRNA and Transcription Factor Gene Regulatory Network Analysis Reveals Key Regulatory Elements Associated with Prostate Cancer Progression. PloS one, 11, e0168760.

49. Lin, Y., Zhang, Q., Zhang, H.M., et al. (2015) Transcription factor and miRNA co-regulatory network reveals shared and specific regulators in the development of B cell and T cell. Scientific reports, 5, 15215.

50. Safran, M., Dalah, I., Alexander, J., et al. (2010) GeneCards Version 3: the human gene integrator. Database: the journal of biological databases and curation, 2010, baq020.

51. Leinonen, R., Sugawara, H., Shumway, M., et al. (2011) The sequence read archive. Nucleic acids research, 39, D19–21.

52. Gupta, A., Ragumani, S., Sharma, Y.K., et al. (2019) Analysis of Hypoxiamir-Gene Regulatory Network Identifies Critical MiRNAs Influencing Cell-Cycle Regulation Under Hypoxic Conditions. MicroRNA, 8, 223–236.

53. Wang, R., Zhang, P., Li, J., et al. (2016) Ubiquitination is absolutely required for the degradation of hypoxia-inducible factor--1 alpha protein in hypoxic conditions. Biochemical and biophysical research communications, 470, 117–122.

54. Dengler, V.L., Galbraith, M., Espinosa, J.M. (2014) Transcriptional regulation by hypoxia inducible factors. Critical reviews in biochemistry and molecular biology, 49, 1–15.

55. Souvannakitti, D., Peerapen, P., Thongboonkerd, V. (2017) Hypobaric hypoxia down-regulated junctional protein complex: Implications to vascular leakage. Cell adhesion & migration, 11, 360–366.

56. Dunnwald, T., Gatterer, H., Faulhaber, M., et al. (2019) Body Composition and Body Weight Changes at Different Altitude Levels: A Systematic Review and Meta-Analysis. Frontiers in physiology, 10, 430.

57. Hu, Q., Ma, T., Wang, K., et al. (2012) The Yak genome database: an integrative database for studying yak biology and high-altitude adaption. BMC genomics, 13, 600.

