## Supplementary for "HAHmiR.DB: A Server Platform For High Altitude Human miRNA-Gene Coregulatory Networks And Associated Regulatory-Circuits"

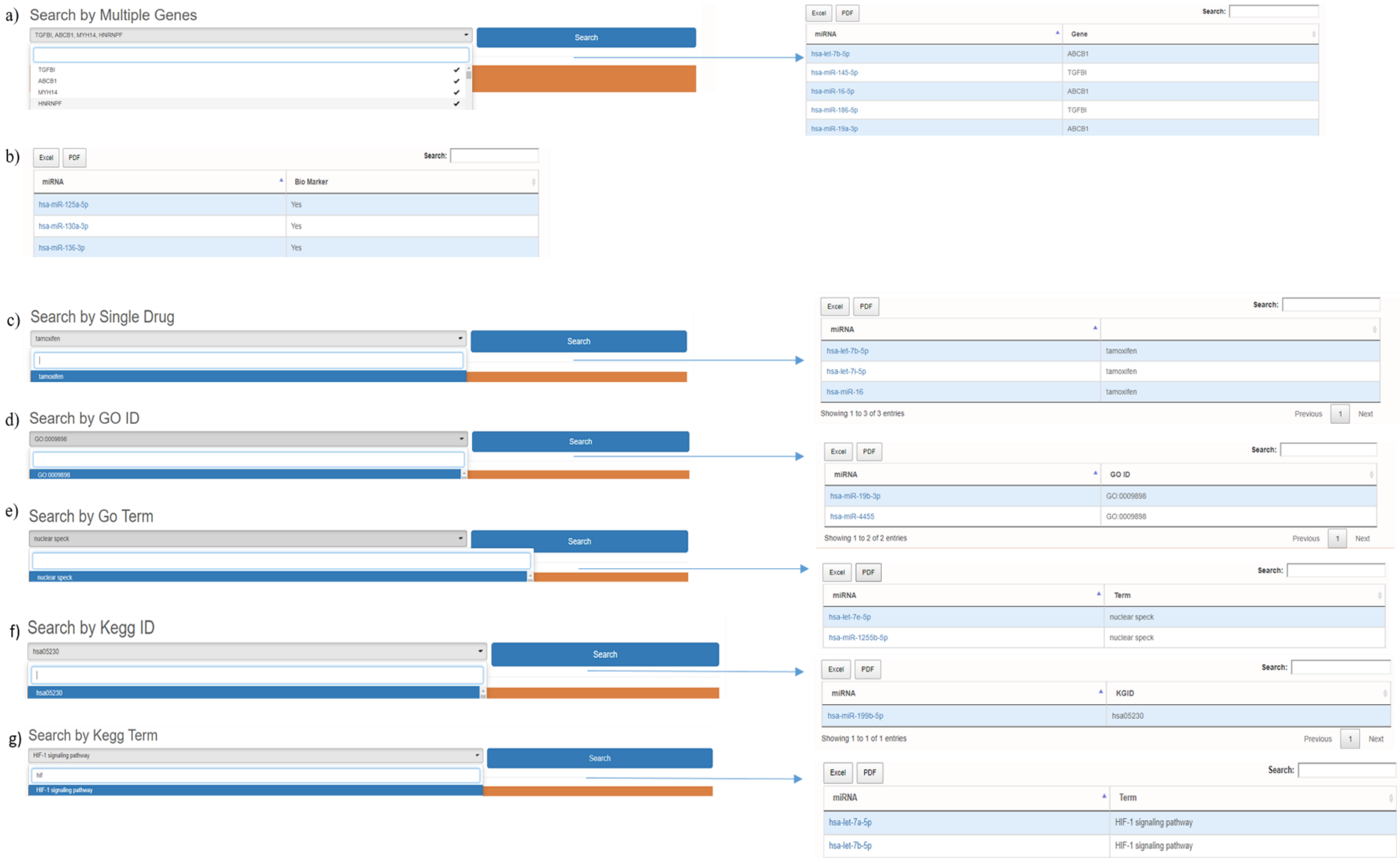


*Figure S1: The web images shows the 'browse by association' option. User can search miRNAs based on a) user defined gene list, b) association as biomarker, c) association with drug d) association with GO ID e) association with GO term f) association with and KEGG ID or name g) association with KEGG pathway name.*
